# Recombination between co-infecting herpesviruses occurs where replication compartments coalesce

**DOI:** 10.1101/345918

**Authors:** Enosh Tomer, Efrat M. Cohen, Nir Drayman, Amichay Afriat, Matthew D. Weitzman, Assaf Zaritsky, Oren Kobiler

**Affiliations:** Department of Clinical Microbiology and Immunology, Sackler School of Medicine, Tel Aviv University, Tel Aviv, Israel; Institute for Genomics and Systems Biology and Institute for Molecular Engineering, University of Chicago, Chicago, IL, USA; Department of Pathology and Laboratory Medicine, Perelman School of Medicine, University of Pennsylvania, Philadelphia, PA, USA; Division of Protective Immunity, Children’s Hospital of Philadelphia, Philadelphia, PA, USA; Department of Software and Information Systems Engineering, Ben-Gurion University of the Negev, Be’er Sheva, Israel.

**Author notes:** Corresponding author: Oren Kobiler.

## Abstract

Homologous recombination (HR) is considered a major driving force of evolution since it generates and expands genetic diversity. Evidence of HR between co-infecting herpesvirus DNA genomes can be found frequently, both *in vitro* and in clinical isolates. Each herpes simplex virus type 1 (HSV-1) replication compartment (RC) derives from a single incoming genome and maintains a specific territory within the nucleus. This raises intriguing questions about where and when co-infecting viral genomes interact. To study the spatiotemporal requirements for inter-genomic recombination, we developed an assay with dual-color fluorescence *in situ* hybridization which enables detection of HR between different pairs of co-infecting HSV-1 genomes. Our results revealed that when viral RCs enlarge towards each other, there is detectable overlap between territories of genomes from each virus. Infection with paired viruses that allow visualization of HR correlates with increased overlap of RCs. Further, inhibition of RC movement reduces the rate of HR events among co-infecting viruses. Taken together, these findings suggest that inter-genomic HR events take place during replication of HSV-1 DNA and are mainly confined to the periphery of RCs when they coalesce. Our observations have implications on understanding the recombination restrictions of other DNA viruses and cellular DNA.

## Introduction

Recombination is considered to be a major driving force in evolution of most organisms, since it accelerates adaptation (1,2). The architecture of eukaryotic nuclei is suggested to regulate many DNA-mediated processes, including replication, gene expression and recombination. DNA viruses that replicate inside the nucleus clearly change the nuclear architecture, however they are subjected to similar spatial constraints as host DNA. The rate of mutation accumulation is lower for DNA viruses than that of viruses with RNA genomes (3,4). It has been hypothesised that high rates of recombination can facilitate genetic adaptation to the changing environment (5). Indeed, homologous recombination (HR) among co-infecting Herpes simplex virus type 1 (HSV-1) genomes is very frequently observed in both *in vitro* genetic assays (6–12) and in sequence analysis of clinical isolates (13–15). Herpesvirus infection therefore provide a system to study spatial features that promote or constrain recombination in the eukaryotic nucleus.

Like all other herpesviruses, HSV-1 viral gene expression, replication and capsid assembly all occur in the host nucleus of infected cells. Viral genomes enter the nucleus through the nuclear pore complex as naked DNA molecules (16), and these rapidly recruit several host and viral proteins to the viral genomes (17–25). Expression of the immediate early and early viral genes allows initiation of viral DNA replication (26). HSV-1 DNA replication proceeds at distinct foci within the nucleus known as replication compartments (RCs) (27,28). The formation of the viral RCs was suggested to initiate from small pre-RCs (29,30). Live cell imaging of viral DNA binding proteins suggested that the pre-RCs migrate toward nuclear speckles, sites of RNA processing, and come into contact with other pre-RCs, where they seem to coalesce into large mature RCs (31). On the other hand, direct visualization of the viral DNA suggested that each RC usually emerges from a single incoming genome (32,33). Our previous study with the swine alphaherpesvirus Pseudorabies virus (PRV) suggested that although viral replication compartments are found in close proximity, they retain distinct territories for each individual genome (33). Earlier experiments with HSV replicons also supported this notion (32). A recent study showed that viral genomes entering the nucleus are observed as condensed foci, and suggested that viral expression and DNA replication allow decondensation of these genomes and formation of RCs (34). Interestingly, some genomes remain highly condensed at the edge of newly developing RCs (34,35). Here, we visualised co-infecting HSV-1 genomes and confirmed that alphaherpesviruses RCs initiate from single genomes.

Viral DNA recombination is facilitated by both viral and cellular proteins. Two viral proteins have been suggested to work as a complex to facilitate viral recombination and have been shown to catalyze strand exchange *in vitro*: the single strand binding protein ICP8 and an exonuclease UL12 (36). Single strand annealing was found to be a recombination mechanism upregulated during viral infection and thus is considered as the mechanism by which the viral recombinase induces recombination (37). While ICP8 is required for viral DNA replication, UL12 is not essential for DNA replication *per se*, although it is required for formation of infectious viral genomes that can be packaged into capsids (38). Recent observations suggest a complex relationship between HSV and the host DNA repair machinery. Some components of the DNA damage response (DDR) machinery are recruited to replicating viral DNA, where they may function in ways that are beneficial for viral progeny (12,21,23,39–42). In contrast, DDR pathways as a whole may be considered to be anti-viral and are suppressed in HSV-infected cells (19,43–48). These findings suggest that the processes of viral recombination and viral replication are intimately associated (46). While knowledge regarding the molecular aspects of HR has been accumulating over the last few years, little is known regarding spatiotemporal constraints on inter-genomic recombination.

The compartmentalization of co-infecting genomes at different RCs raises questions about where and when recombination takes place during the course of infection. To tackle these questions, we utilized a fluorescence *in situ* hybridization (FISH) based assay to differentiate between *de-novo* synthesized variants of viral genomes. Our results suggest that multiple inter-genomic recombination events occur at later stages of infection following DNA replication, and that inter-genomic recombination takes place at the interface between mature RCs. We also found that the number of RCs correlates with nuclear size, suggesting a possible spatial restriction to the number of viral genomes that initiate replication.

## Materials and methods

### Cell culture

African green monkey kidney cells (Vero, ATCC CCL-81) and human female osteosarcoma cells (U2OS cells ATCC HTB-96) were grown in Dulbecco’s Modified Eagle Medium (DMEM X1; Gibco), supplemented with 10% Fetal Bovine Serum (FBS; Gibco) and 1% Penicillin (10,000 units/ml) and Streptomycin (10 mg/ml; Biological Industries, Israel).

### Viruses

All viral recombinants are derivatives of herpes simplex type 1 strain 17. Each viral recombinant contains one or two tag sequences for specific staining by FISH. To facilitate isolation, both tag sequences are expression constructs for fluorescent proteins. The red fluorescent protein mCherry driven by the human cytomegalovirus promoter (CMVp) and the yellow fluorescent protein YPET driven by the simian virus 40 promoter (SV40p). Tag sequences were inserted into the viral genome by homologous recombination. Viral DNA was co-transfected along with a plasmid containing the tag sequences flanked by sequence homologies to the viral site of insertion (synthetically generated by GenScript, Piscataway, NJ, USA). Recombinant viruses were isolated from the progeny by plating lysate from transfected Vero cells and picking fluorescent plaques using an epi-fluorescent microscope. Viral stocks were prepared by growing purified plaques for each recombinant virus on Vero cells. The viral recombinants were validated by PCR. Viral titers were measured by plaque assay. An additional viral recombinant containing both tag sequences was isolated by crossing the recombinant OK26 to the previously described OK11 (49) and selecting for plaques containing two fluorescent proteins by plaque assay. All viral recombinants constructed for this paper are described in Table 1

**Table 1.**
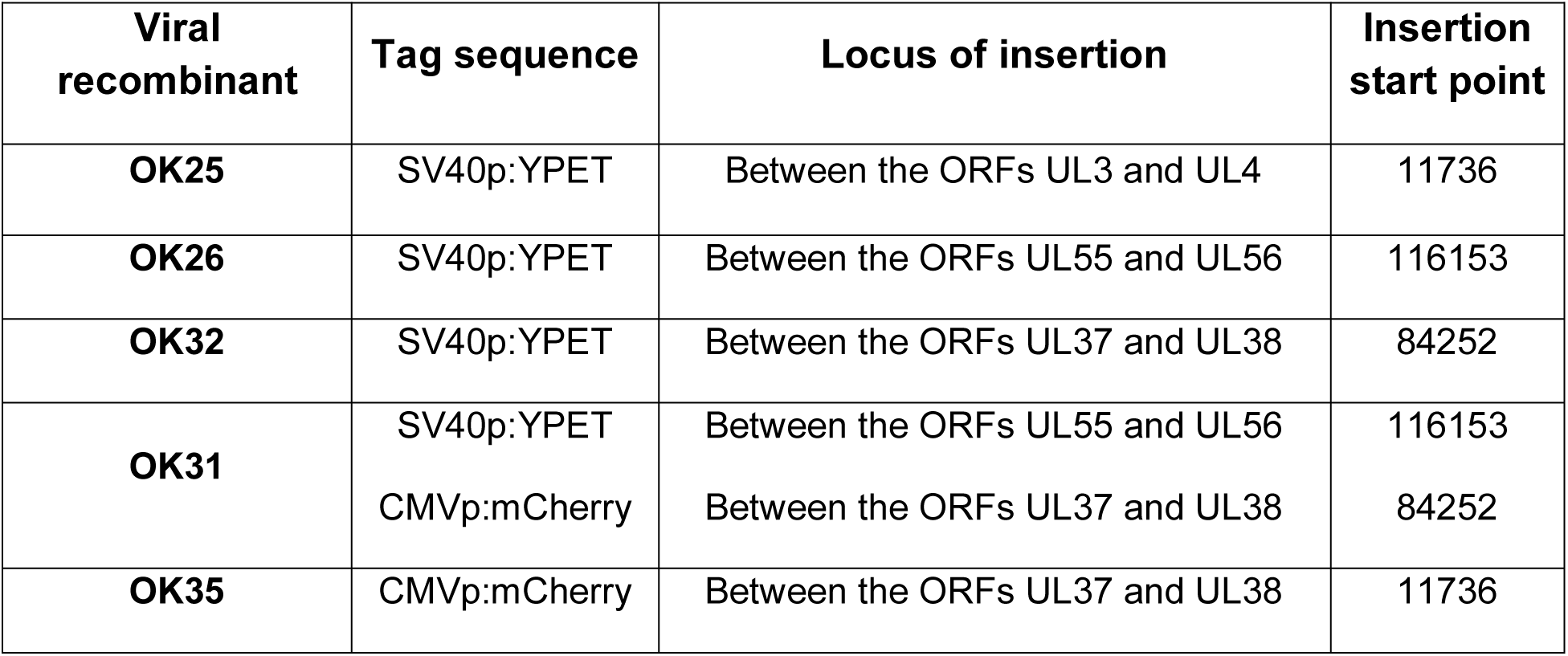
List of viral recombinants used in this work.

### Fluorescent probes

A set of 20 fluorescent probes was designed to correspond to each one of the two tag sequences. The probes for the CMVp:mCherry and sequences SV40p:YPET were conjugated on their 5' end to the fluorescent dyes cy3 and cy5 respectively. An additional probe was designed to stain HSV1 viral DNA non-specifically. This probe corresponds to the viral a’ sequence and conjugated to the fluorescent dye Alexa Fluor 488 on its 5’ end. Fluorescent probes were synthesized by Integrated DNA Technologies (Coralville, IA, USA). The probes were dissolved in TE buffer to a stock concentration of 10 μM. Probes from each set were pooled together in equal ratios and kept in −20°C before hybridization. Probes sequences are detailed in Table 2.

**Table 2.**
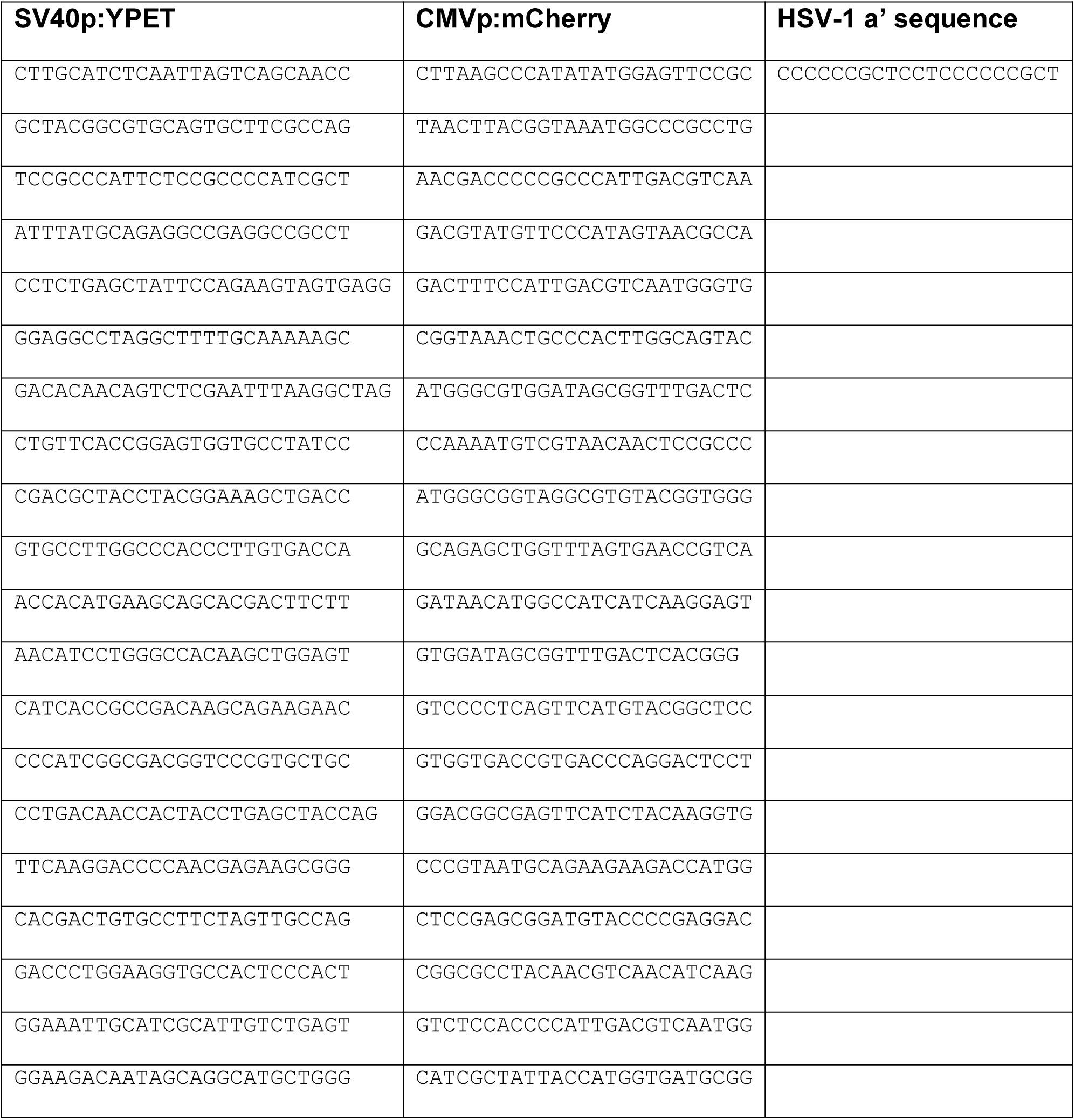
List of fluorescently labelled probes for FISH.

### Fluorescent in situ hybridization

Cells where seeded and grown to confluence inside 12 well plates underlined with a glass coverslip. Cells were infected at MOI 20 with 200 µl inoculums. The cells were incubated on ice for 1 hour following the addition of the inoculums. The inoculums were then removed and replaced with DMEM+10%FBS. The infected cells were incubated on 37°C for 6 hours. Medium was then removed and cells were washed with PBS. Cells were then fixed with 1 ml 4% Paraformaldehyde (PFA) in PBSX0.3 for 10 minutes at room temperature. Cells were washed three times with 0.05% Triton X-100 in PBS at 5-minutes intervals. Cell nuclei were permeabilized by incubating the coverslips with 0.5% Triton X-100 in PBS for 20 minutes. The 0.5% Triton was then removed and 20% glycerol (Sigma-Aldrich) in PBS was added and incubated at room temperature for 30-60 minutes. Following incubation, coverslips were repeatedly frozen and thawed by dipping them in liquid nitrogen three times. Before refreezing, the coverslips were dipped in the 20% glycerol solution to prevent cells from drying. Coverslips were then washed with 0.05% Triton X-100 in PBS 3×5 minutes. Coverslips were briefly washed in 0.1N HCl and incubated in fresh 0.1N HCl for 8 minutes. Subsequently, the coverslips were washed with 0.05% Triton X-100 in PBS 3×5 minutes and then transferred to 50% formamide (Merck) in Saline Sodium Citrate buffer X2 (SSCX2; diluted from SSCX20 prepared as follows: 175.3gr Sodium Chloride (Bio-Lab), 88.2gr of Sodium Citrate (Merck) in 1 L of DDW, adjusted to pH 7.0 and filtrated). Viral genomes were hybridized using hybridization solution (10% Dextran Sulfate (Calibochem) in SSCX2) containing the fluorescent probes (IDT; final volume of 0.7 uM for each probe). For each coverslip, 5 µl of probe was placed on a clean slide. Each slide was lifted out of the 50% formamide in SSCX2 solution and coverslips were placed directly on the probe drop, cell side down and excess fluid was removed. Rubber cement was applied to the edges of the coverslip to form a temporary seal for hybridization. The slides were set aside to dry completely at room temperature. The denaturing of cellular and probe DNA was achieved by placing the slides on a hot slide block (Eppendorf) for precisely 2 minutes at 95°C. Following denaturation, slides were sealed with parafilm and placed in a 37°C incubator for 72 hours. After sufficient hybridization, the rubber cement was removed from the slides. Each coverslip was carefully removed and washed with SSCX2 3×5 minutes at 37°C then washed with SSCX0.1 3×5 minutes at 63°C. For each coverslip, one drop of Fluoroshield mounting medium containing DAPI (Abcam, Cambridge, MA, USA) was placed on a fresh slide and each coverslip was placed cell side down on the drop. Coverslips were sealed with nail polish. FISH protocol was adapted from (50).

### Microscopy

Viral plaques were visualized under a Nikon Eclipse Ti-E epifluorescence inverted microscope. Single cell analysis by FISH assay was performed using a Nikon Eclipse Ti microscope equipped with Yokogawa CSU X-1 spinning disc confocal system.

### Image analysis

Nuclei segmentation: Nuclei were automatically segmented from the DAPI channel as follows. Gaussian smoothing (**σ** = 2) was applied to the image followed by Otsu thresholding (51) to partition the image to nuclei / background regions. The nuclei segmentation was refined by morphological operators: opening (width = 3 pixels) to exclude small excesses; closing and an additional opening (width = 20 pixels) to unify regions that belong to the same nucleus; Filling holes; considering only connected regions of area exceeding 900 pixels - the minimal nucleus size. The output of this stage is a set of masks, each corresponds to a single nucleus.

RC segmentation: RCs were segmented from each fluorescent channel independently. Three groups of intensities were observed within each nucleus: background - at intensities close to those outside the nucleus; intermediate - above the background levels; and bright - which are the replication centers. First, Gaussian smoothing was applied (**σ** = 2) to the image. Second, background statistics were calculated (mean intensity, standard deviation) from the pixels outside the nuclei masks. A threshold of 2.5 standard deviations above the background mean intensity level was calculated. To exclude pixels with background intensities from further analysis, the threshold was applied for each nucleus mask. Otsu thresholding was then applied to the remaining pixels, pixels below the threshold were pooled and a new threshold of two standard deviations above the mean was calculated. The pixels above these thresholds were defined as the RCs. The segmentation refined by applying morphological operators: opening - excluding small excess, closing - filling gaps, and opening again - disconnecting independent replication centers; all with a square kernel of 3 pixels width. Last, were holes filled to define the RCs. Parameters were optimized by visual assessment of RC segmentation accuracy.

Manual selection of nuclei: Region for statistical analysis were manually selected based on the accurate segmentation of RCs. Regions of accurate RC segmentation were selected independently for each channel, the intersection (implying accurate segmentation on both channels) was used for quantifications.

The program outputs required measurements such as nuclei area, number and area of RCs in each channel of each nucleus and the size of overlapping area for each RC. The data were then transferred into Microsoft excel for subsequent processing. The data were collected for each nucleus for further analysis. Cells without colocalization were removed from further analysis. The ratio of colocalization area out of the total area of RCs was calculated per cell (see supplementary excel file). Outliers were removed using the interquartile method (1.5 times interquartile distance from either the 1^st^ of 3^rd^ quartile). Two tails students T.test was performed to determine significance.

Code availability: The Matlab source code used for analysis is publically available at https://github.com/assafzar/RecombinationHSV1.

### Fluorescent plaque assay

Cells (either Vero or U2OS) were grown to confluence in 12 well plates. Infection was carried out at an MOI of 20 with a mix of the co-infecting viruses in 4°C. Cells were harvested 9HPI. In Latrunculin B experiments, 1HPI the medium was replaced by medium containing 2.5 μM Latrunculin B (Sigma). The infected cells were lysed by freeze thawing and sonicating. The lysate was plated on 6 well plates containing Vero cells. Following infection, cells were overlaid with growth medium containing 0.5% methylcellulose and incubated for 48 hours. The plates were then inspected under fluorescent microscope for plaque counting. The percent of dual color plaques out of the total fluorescent plaques are indicated. A total of 8 wells, collected from two individual plaque plating, from two technical repeats in two separate experiments, were obtained. Modified Thompson Tau Test was used for outlier removal (no more than one outlier per condition was detected). Two tails students T.test was performed to determine significance.

## Results

### A FISH based assay designed to detect inter-genomic recombination events between co-infecting viruses

Recombination among co-infecting HSV-1 strains is a frequent event that can be detected by the progeny viruses released from infected cells (8–10,12). To study the spatiotemporal constraints of these inter-genomic recombination events we developed a FISH based experimental assay that enables visualization of two different viral genomes within a co-infected nucleus. For this assay, we constructed a series of viral isolates, isogenic to each other except for two unique tag sequences. These tag sequences, YPET or mCherry encoding genes (yellow or red fluorescent proteins, respectively) were inserted into various loci across the viral genome (Figure 1A and Table 1). We designed two sets of fluorescent probes, one set for each tag sequence (Figure 1B). Each probe set was conjugated to a distinct fluorophore (Cy3 or Cy5) to enable visual identification of the genomes containing each tag sequence. We hypothesize that using different mixtures of viruses should lead to distinct patterns within the infected nucleus (Figure 1C). As was shown previously for PRV (33), we expect that co-infection with two HSV-1 viruses containing tag sequences at the same genomic locus cannot result in a new recombinant genome containing both tags. Thus, each RC will react to a single fluorophore, including at the contacting edges of proximate RCs (Figure 1C example I). Infection with a viral recombinant containing two tag sequences within one genome is expected to result in RCs stained with both fluorophores (dually-labelled RCs) (Figure 1C example IV). Infection with two viral recombinants containing tag sequences at different genomic loci could result in progeny genomes that either contain both tag sequences on a single genome or contain neither tag sequence. One of two types of spatial patterns could be expected to dominate under these conditions. Dually-labelled RCs (fully covered by both probe sets) would imply that inter-genomic recombination takes place early during infection before the viral DNA replicates and RCs mature. In this situation, the reciprocal recombinant genome would contain no tag sequences and will generate RCs that are not detected since they are not covered by any of the probes (Figure 1C example II). Alternatively, partially overlapping RCs that fuse to each other at their periphery, would suggest that inter-genomic recombination occurs later during the infection cycle following viral DNA replication (Figure 1C example III). Therefore, this FISH assay is designed to detect viral inter-genomic recombination events, with visual readouts for when and where it takes place.

**Figure 1.**
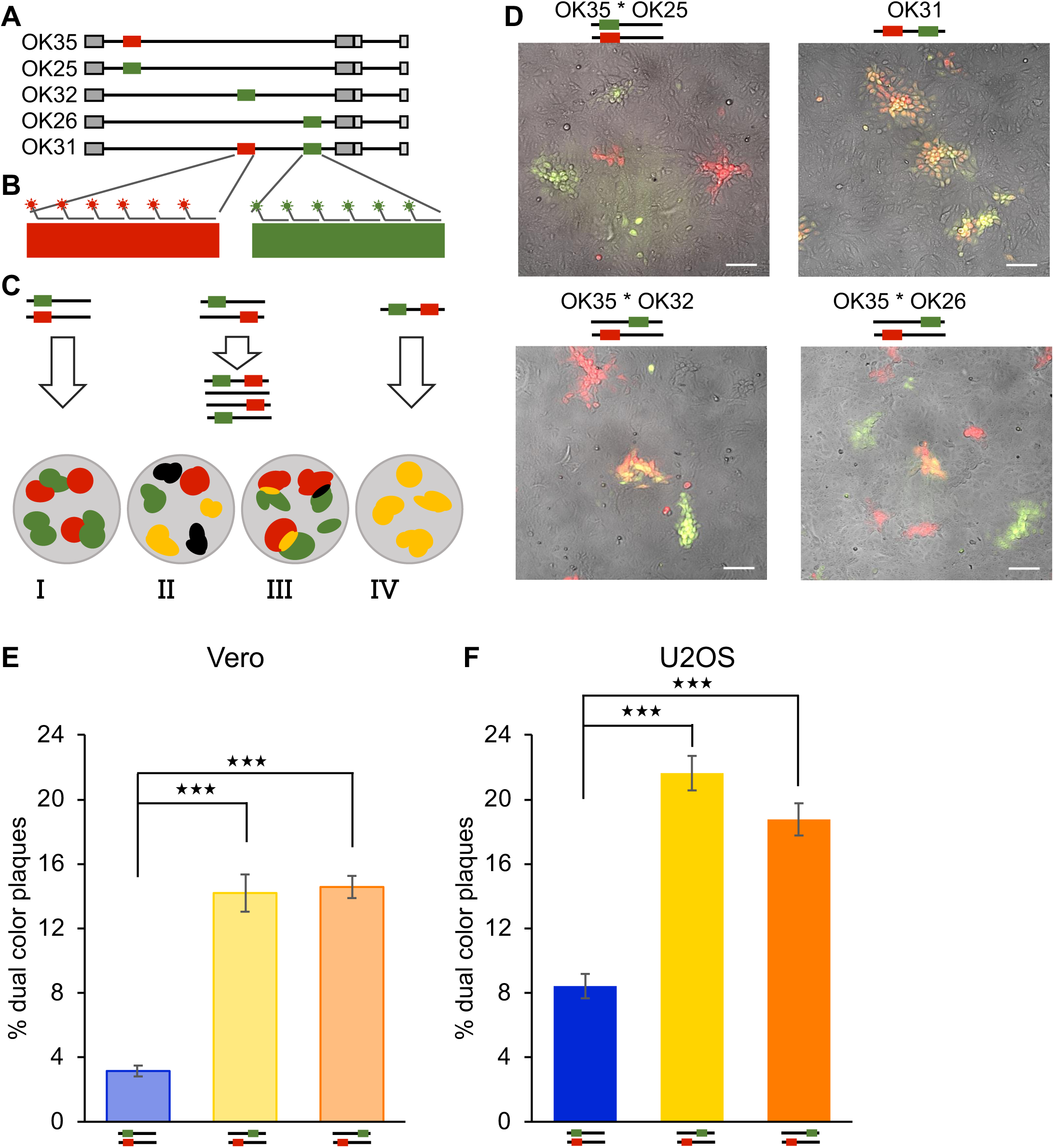
Schematic representation of dual colour FISH assay. **(A)** A series of viral recombinants with two unique tag sequences (designated red and green) inserted into various loci of the parental genome. Illustration of the insertion sequences (colour boxes) and viral genomes (black line with repeats marked in gray boxes). Illustrations are not to scale. **(B)** Two sets of probes conjugated to distinct fluorophores, each set corresponds to one tag sequence. **(C)** Illustration of expected results, in which the gray nucleus contains viral RCs originating from genomes with either one tag sequences (red or green), two tag sequences (yellow) or no tag sequence (black). The expected results in the case of infection with tag sequences on the same locus of two co-infecting genomes **(I)**, with tag sequences on separate loci in two co-infected genomes where recombination takes place before DNA replication **(II)** or following DNA replication **(III)** and with two tag sequences in the same genome **(IV)**. **(D)** Representative images of viral plaques initiating from progeny viruses collected from Vero cells co-infected with different viral recombinants as marked above each image. Scale bar is 100 µm. **(E-F)** Progeny viruses from either Vero **(E)** or U2OS **(F)** cells co-infected with the viral recombinants OK35 & OK25 (blue), OK35 & OK32 (yellow) and OK35 & OK26 (orange) were plated for individual plaques and colors quantitated. The percent of dual color plaques are indicated. The averages are shown from two individual plaque plating, from two technical repeats in two separate experiments. Error bars represent standard error of the means. ***P < 0.001; by t test.

### Patterns of RC interactions dependent on HR

To identify inter-genomic recombination events, we co-infected Vero cells with different pairs of viruses listed in Table 1, as detailed below. All infections were carried out at MOI 20, to increase the likelihood of interactions between incoming viral genomes (8). Cells were either infected with one virus that contained both mCherry and YPET tag sequences (OK31) or were co-infected with two viruses, each carrying a different tag sequence (OK35 together with either OK25, OK32 or OK26). First, we collected the progeny viruses released from the infected cells at 9 hours post infection (HPI) and plated them for single plaques. We found that co-infections can result in dual color (yellow) plaques under our different infection conditions (Figure 1D). The number of dual color plaques was measured for each infection pair and compared to the total number of plaques (Figure 1E). We found that the rate of dual color plaques was increased by ~5 fold for infections with viral genomes carrying fluorescent protein genes at different genomic sites when compared to infections with viruses where the genes are located at the same genomic site. We note that there was no statistical difference between the two infections with the viruses carrying the reporter genes at alternative sites in the genomes (OK35 together with either OK32 or OK26). Our results corroborate the conclusion that most progeny viruses containing dual color are the outcome of inter-genomic HR events (8).

To visualize inter-genomic HR events at the single cell level, cells were co-infected and fixed at 6HPI for hybridization with the appropriate fluorescent probes. Using confocal fluorescent microscopy, we imaged the viral RCs and identified four distinct patterns of interactions between the RCs (Figure 2A-D). The first interaction type (A - minimal overlap) is of two RCs that come into close contact but without evidence of mixing between the two RCs, i.e. no visible co-localized pixels and no intersection between RCs margins (Figure 2A). These interactions were observed in all co-infections, however they are significantly more pronounced when the tag sequences were at the same relative genomic locus (OK25 and OK35, P<0.05, Figure 2E). The second interaction type (B - periphery overlap) is of two RCs that come into close contact and clearly mix with each other, as defined by the presence of dual color pixels at the site of contact (Figure 2B). Note that the two channels do not necessarily co-localize at the single pixel resolution at the contact sites (i.e., no yellow pixels), but do co-localize at the region-scale. This was the most common interaction observed in all co-infections (Figure 2E), with significantly increased frequency when the two tag sequences were located at different sites of the viral genome: either OK26 and OK35 (P<0.005) or OK32 and OK35 (P<0.05). In the OK25 and OK35 dual infection (tags located at the same genomic locus) this pattern cannot be due to HR, and in this case reflects limitations of image resolution. In the third interaction type (C-Full overlap within larger RC), a small RC (smaller than half the size of all other RCs in that nucleus) is fully contained in a much larger RC (Figure 2C). This pattern was detected in about 5% of all co-infections (Figure 2E). The fourth interaction type (D – Full overlap) is a dual color RC, in which the entire area of the RC contains both fluorophores (Figure 2D). The relative frequencies of these events can be quantitated during single or dual infections (Figure 2E). As expected, the overlapping pattern accounted for ~ 90% of RCs in cells with single infection of virus carrying both tag sequences (OK31), and in about 3% of interactions in all dual co-infections. We deduce that our assay can be used to readout inter-genomic recombination, since detectable HR events change the overall distribution of the interactions among the RCs. In all co-infection assays, partial overlaps at the point of contact are more frequent, compared to entirely overlapping RCs. The rare occurrence of entirely overlapping RCs compared to the high frequency of HR during HSV-1 co-infection (12) suggests that these RCs capture infrequent HR events and cannot reflect the majority of HR between RCs. We therefore suggest that inter-genomic HR occur at points of physical interaction between co-infecting genomes, after viral DNA replication has initiated.

**Figure 2.**
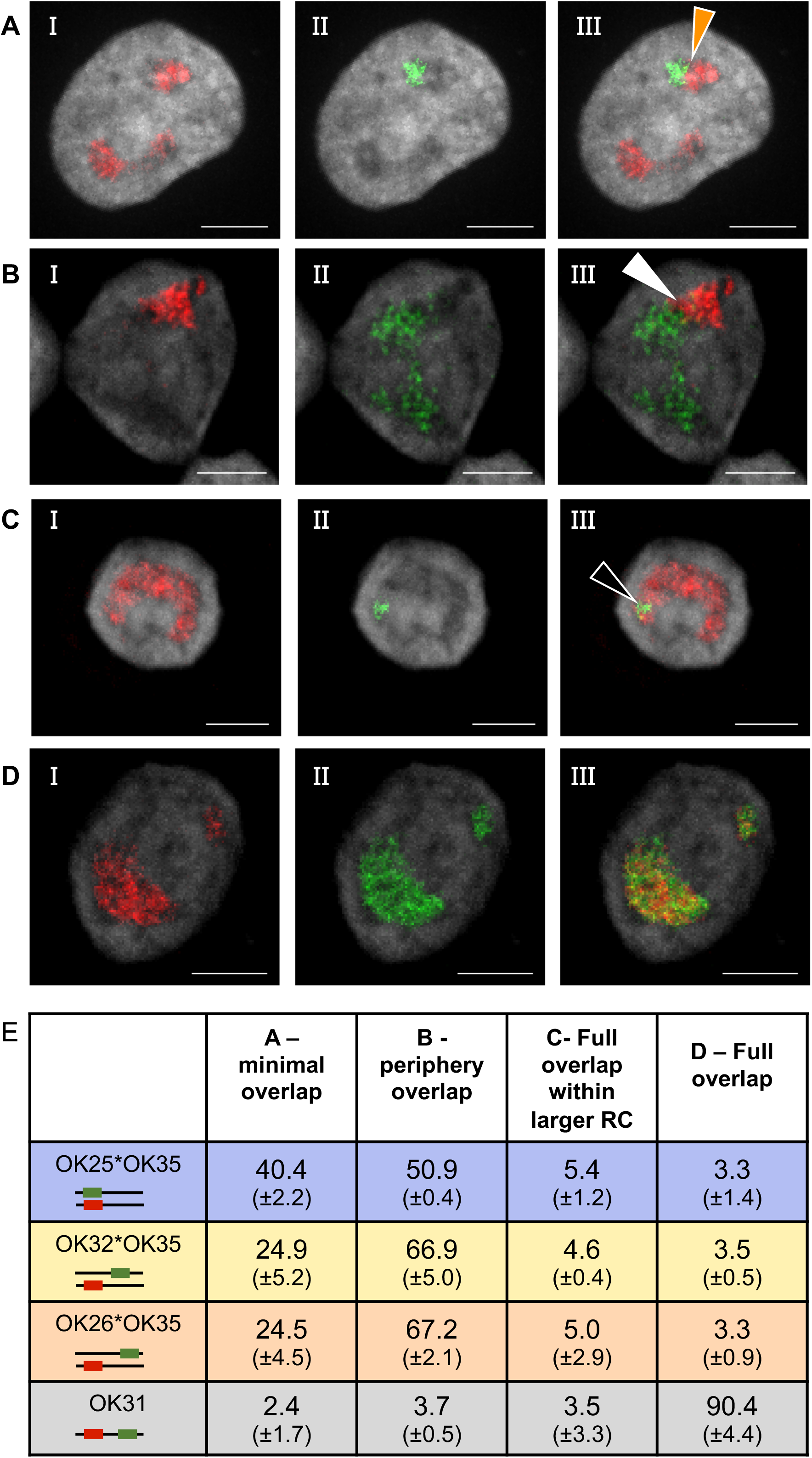
Observed patterns of RC interactions. **(A-D)** Representative images of Vero cells infected with viral isolates containing two genomic tags, each corresponding to a set of fluorescent probes. The probe sets are imaged separately in red and green (I and II) and merged (III). Four observed patterns of interaction between RCs are represented: **(A)** RCs containing different tag sequences come into contact without overlap (orange arrowhead), imaged from cells infected by two viral recombinants containing tag sequences on the same genomic locus (OK25 and OK35). **(B)** RCs containing different tag sequences overlap at their periphery (white arrowhead), imaged from cells infected with two viral recombinants containing tag sequences on separate genomic loci (OK32 and OK35). **(C)** RC containing one tag sequence overlap completely within a larger RC with the second tag (black arrowhead) imaged from cells infected with two viral recombinants containing tag sequences on separate genomic loci (OK26 and OK35). **(D)** Entirely overlapping RCs, imaged from cells infected by a viral recombinant containing two tag sequences on the same genome in separate loci (OK31). DAPI nuclear stain is presented in gray. Scale bar is 5µm. **(E)** Percent of each of the observed RCs interactions (A to D) from the total number of RCs interactions. Manually counted at the different co-infection conditions. >70 RCs interactions per infection type per experiment were counted. An average of three experiments and the standard deviation in brackets are shown.

We have previously shown that infections of U2OS cells result in more HSV-1 genomes initiating gene expression per cell when compared to infections of Vero cells (52). To test if the number of initiating genomes has an effect on the interactions between RCs, we repeated our dual color infection assay in U2OS cells. Although overall higher levels of dual-colored plaques were observed in U2OS cells when compared to Vero cells, the relative levels of dual colored plaques attributed to HR (i.e co-infection with either OK26 and OK35 or OK32 and OK35 relative to co-infection with OK25 and OK35) was not different between the two cell types (Figure 1F). In both U2OS and Vero cells there was no statistical difference between HR rates when comparing infections with viruses that carry the second reporter gene at different locations in the viral genome (OK26 versus OK32).

In FISH based experiments with infections of U2OS cells, we observed all four patterns of RCs interactions that we defined for infections of Vero cells (Figure 3). However, in U2OS cells, multiple RC interaction patterns in a single cell were more common than in Vero cells (Figure 3A and 3B), probably due to the higher number of RCs observed per cell in this cell type (see below, Figure 5A). The distribution of patterns observed for the different infections in U2OS cells (Figure 3D) was similar to the distribution detected in Vero cells (Figure 2E) although in U2OS cells lower rates of type C and D interactions were observed in all co-infections tested.

**Figure 3.**
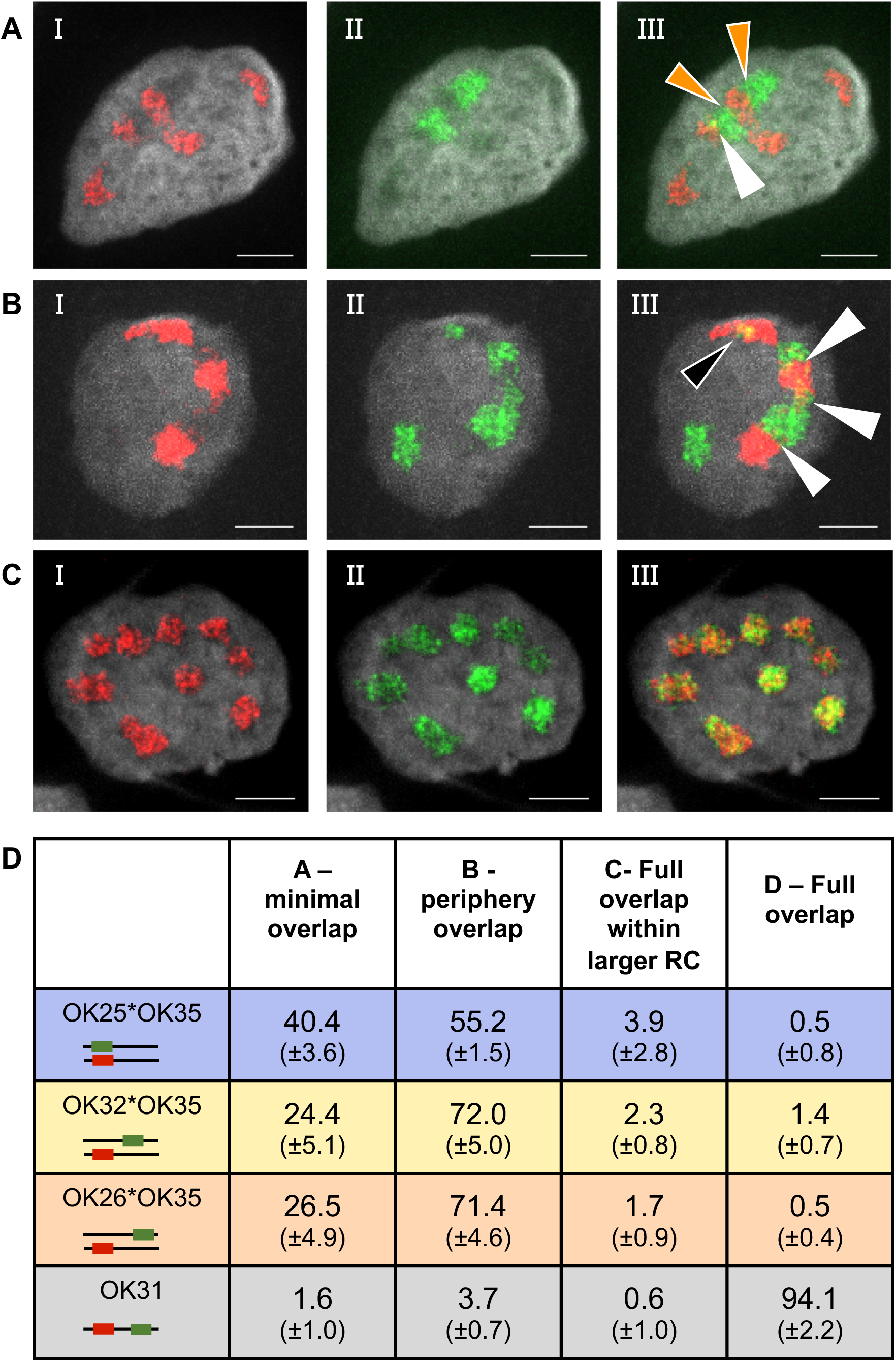
Patterns of RC interactions on U2OS cells. **(A-C)** Representative images of U2OS cells infected with either **(A)** OK25 and OK35, **(B)** OK32 and OK35 or **(C)** OK31 viral recombinants. The probe sets are imaged separately in red and green (I and II) and merged (III). Arrowheads are colour coded as in figure 2. DAPI is presented in gray. Scale bar is 5µm. **(D)** Percent of each of the observed RCs interactions out of total RCs interactions. Manually counted at the different co-infection conditions. >70 RCs interactions per infection type per experiment were counted. An average of three experiments and the standard deviation in brackets are shown.

### Overlap between RCs is enhanced by HR

Merely identifying the patterns of overlap does not estimate the relative proportion of the overlapping area between RCs. Therefore, we developed a quantitative evaluation of the RC overlapping areas in each cell. We found that common pixel based co-localization methods do not provide an accurate estimate of the degree of overlap between HSV-1 RCs, probably due to the relaxed form of the replicating viral DNA (34). Our results following infection with one virus carrying the two tag sequences (OK31) showed limited co-localization at the pixel level, similar to results obtained by two probes corresponding to adjacent loci on HSV-1 genomes (53). We therefore applied a semi-automated object-based method to quantify the degree of overlap for RCs in cells infected with viruses that have two distinct genomes (see Methods). For each cell nucleus, we independently segmented the RCs in each channel. The relative overlapping area was calculated as the ratio between the overall area of the overlap and the total overall area occupied by viral DNA (total area of all RCs minus the total overlap area). The images analysed were collected from three infections carried out on different days from separate viral stocks. Over 200 cells from each cell type were analysed for each co-infection. The dual-tagged virus (OK31) allowed us to analyse fewer cells in this infection (N = 120 Vero, N = 99 U2OS) but still with sufficient statistical power.

We analysed all the cells in which contacts between red and green RCs were observed (more than 70% of cells in all co-infections). For both cell types, we found a small but significant increase in the relative overlapping area in each nucleus for cells co-infected by viruses with tags in different viral loci compared to cells co-infected by viruses with tags in the same locus (greater than 16% increase for OK32 and OK35 infection on Vero cells and more than 30% increase in all other co-infections, Figure 4A-B). As with plaque assays (Figure 1E and F) and manual count (Figure 2E and 3D), we did not observe significant differences in overlap between co-infections with viruses that have tags in different loci. To confirm that cellular parameters do not bias our measurements, we compared the relative overlapping area to the total nuclear area (Figure 4C-D) or to the number of RCs per nucleus (not shown). We found no evidence of correlation between these two parameters and the relative overlapping area in all infection conditions (Pearson correlation bellow 0.3 for each of the infections). We conclude that the observed significant increase in overlapping areas indicates the readout of HR events. As mention above, the relative high background levels (OK25 and OK35 infection) of overlapping areas (~20%) are due in part to noise of the assay and to non-HR events during viral replication.

**Figure 4.**
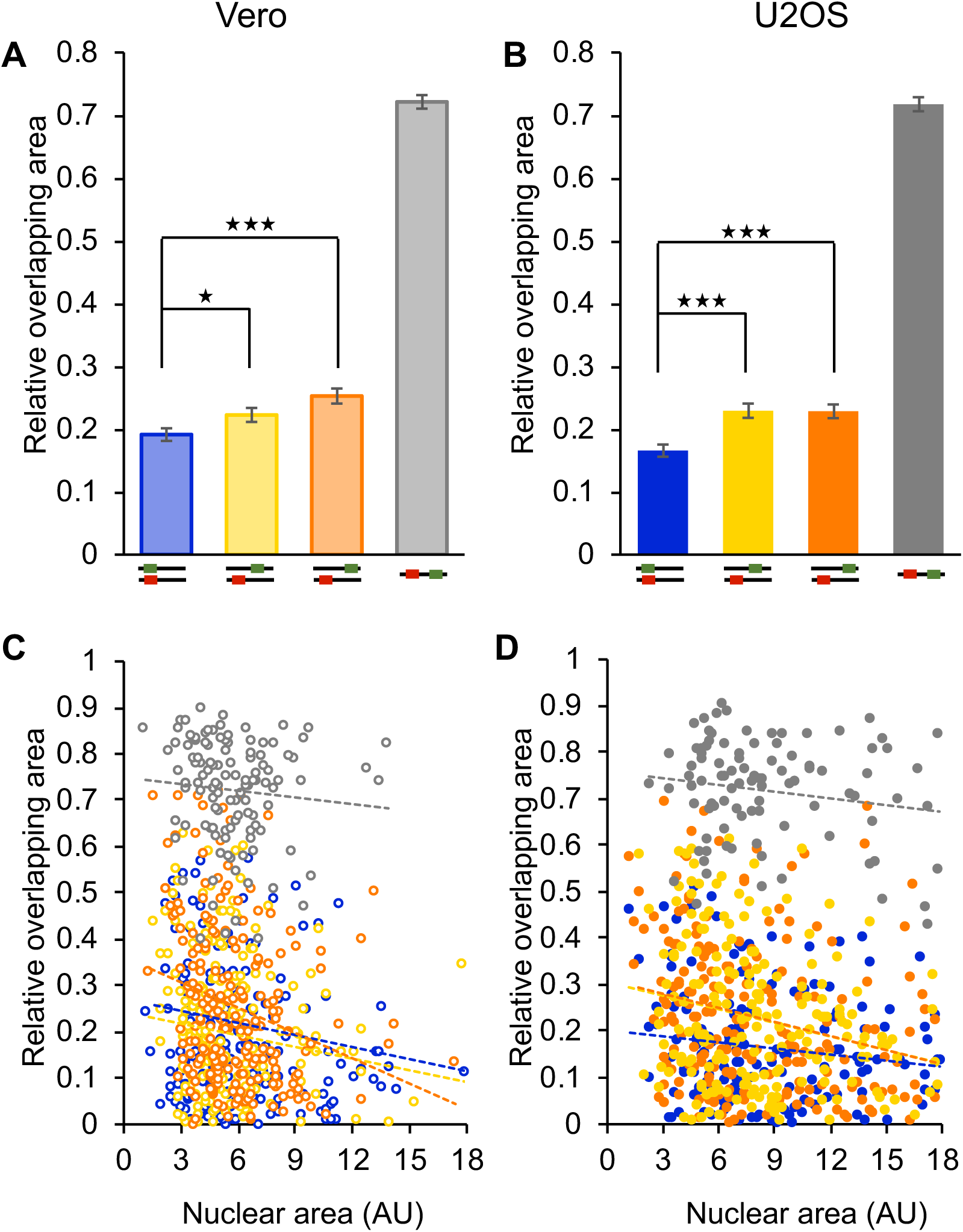
HR enhances overlap between viral genomes. Vero cells **(A, C)** and U2OS cells **(B, D)** were infected with the viral isolates OK35 & OK25 where tags are inserted into the same genomic locus (blue), the viral isolates OK35 & OK32 (yellow) or OK35 & OK26 (orange) containing tags on separate genomic loci and the viral isolate OK31 containing two tag sequences on a single viral genome (gray). **(A and B)** Each column represents the average relative overlapping area calculated from all cells (with detectable overlap) infected under the same condition. Error bars represent standard error of the means. *P < 0.05, ***P < 0.001; by t test. **(C and D)** Individual cells were plotted to compare the relative overlapping area to the nuclear area. All analyses were conducted on images generated from three independent experimental repeats done on different days with viral stocks prepared and tittered separately.

### The number of RCs correlates with nuclear size

We previously demonstrated that the number of HSV-1 genomes replicating per cell can influence the outcome of infection (54) and that host conditions prior to infection impact the outcome of infection at the single cell level (54,55). To test the effect of cellular parameters on RC formation and growth, we took advantage of our assay that provides an opportunity to quantitate the number of mature RCs within individual cells as we can distinguish between the colors of coalescing RCs. We note that OK31 infection was omitted from this analysis due to our inability to distinguish between coalescing RCs in this infection. We therefore tested association of the number of RCs to other parameters of the infected cell. First, we observed that in U2OS cells more RCs were detected per individual nucleus than in Vero cells. We detected an average of ~5.1 RCs per nucleus in U2OS cells compared to ~3.7 RCs per nucleus in Vero cells (1.375 fold increase, Figure 5A). Interestingly, a similar fold increase of ~1.38 was identified in nuclear area (Figure 5B). This led us to speculate that nuclear area may correlate with the number of RCs per nucleus. Indeed, the number of RCs correlates to the nuclear area for individual cells in each of the cell types (Pearson correlation: R > 0.538 for all co-infections, p < 0.0001 for all co-infections, Figure 5C-D). We hypothesize that the increase in the number of RCs per nucleus will result in more total viral DNA (RCs area) in larger nuclei. As expected, the nuclear area correlates with the total area of the RCs per nucleus (Pearson correlation: R = 0.530 for OK32 and OK35 co-infection on Vero and R > 0.625 for all other co-infections, p < 0.0001 for all co-infections, Figure 5E-F). We speculated that the increase in total RCs area could also result from the possibility that RCs expand faster to a larger size in larger nuclei. We therefore compared the mean RC area (per cell) to the total nuclear area (Figure 5G-H). We found a much weaker correlation between these parameters (Pearson correlation: < 0.5 for U20S cells and < 0.3 for Vero cells in each of the infections). These results suggest that the increase in RC area in larger nuclei results from higher numbers of RCs rather than increased RC size. Taken together, our results suggest that nuclear size could be a restricting factor, limiting the number of incoming genomes that are able to initiate replication.

**Figure 5.**
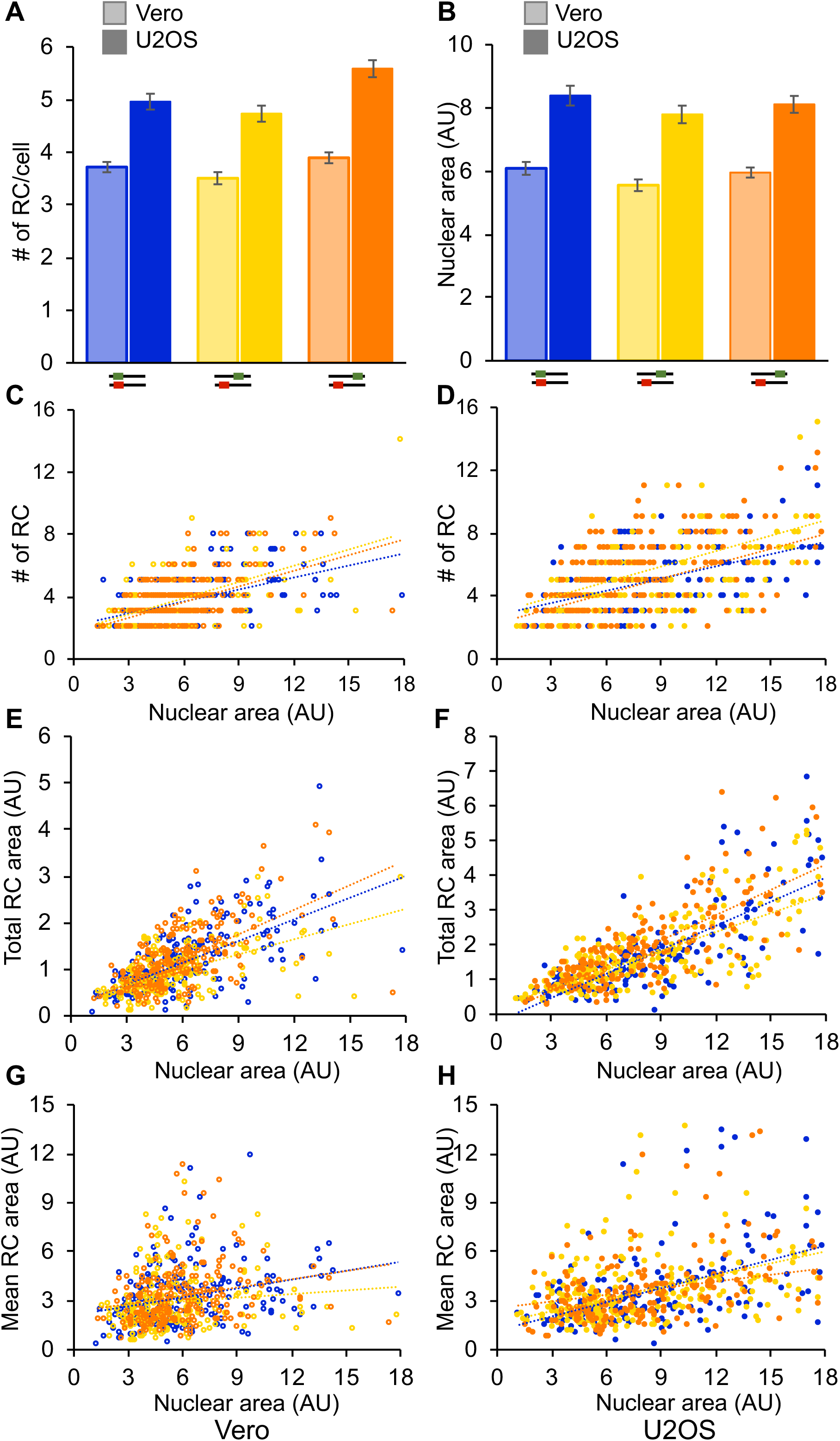
Nuclear size correlates with the number of RCs per cell. Cells were co-infected with the viral recombinants OK35 & OK25 (blue), OK35 & OK32 (yellow) and OK35 & OK26 (orange). Comparing the mean number of RCs per cell **(A)** and the mean nuclear area **(B)** between Vero (bright coloured columns) and U2OS (dark coloured columns). Error bars represent standard error of the means. **(C-H)** Individual Vero **(C, E, G)** and U2OS **(D, F, H)** cells were plotted to compare the number of RCs per cells **(C, D)**, the total area of RCs per cell **(E, F)** and the average area of each RCs **(G, H)** to the nuclear area. A trend line (colour coded as above) was calculated using the ordinary least squares (OLS) method is presented for each infection condition. All analyses were conducted on images generated from three independent experimental repeats done on different days with viral stocks prepared and tittered separately.

### Inhibition of RC movement reduces the rate of HR events

It was previously shown that RC movement towards each other relies upon the formation of actin filaments (31). Since our results suggest that HR events occur between coalescing RCs, we tested whether disruption of RC movement will result in reduction in recombination rates. We treated co-infected cells with latrunculin B, an inhibitor of actin polymerization, and tested for recombination rate among the progeny viruses by fluorescent plaque assay. Vero and U2OS cells were infected with the pairs of viral constructs that produce dual color progeny as a result of HR (OK32 and OK35, OK26 andOK35) and were treated with latrunculin B. At 9HPI, progeny viruses were collected and plated for single plaques. The rate of HR events was compared to the progeny from infection with the pair of viral constructs carrying the fluorescent gene at the same site (OK25 and OK35). We note that treatment with latrunculin B increased the level of dual colored plaques that did not result for HR (from the viral pair OK25 and OK35) compared to untreated cells (Figure 6). To estimate the rate of HR for viral pairs with fluorescent tags in different loci of the viral genome, the average rate of OK25 and OK35 dual colored plaques was subtracted from the total rate of dual colored plaques from the viral pairs OK35 and either OK26 or OK32. The presence of latrunculin B reduced HR rates compared to untreated cells for both infection pairs in both cell types (Figure 6). Statistically significant decreases were observed in both cells during OK26 and OK35 co-infection and only in U2OS cells for OK32 and OK35 co-infection. These results suggest that RC movement enhances inter-genomic HR, corroborating our initial conclusion that HR events occur at the contact point of coalescing RCs.

**Figure 6.**
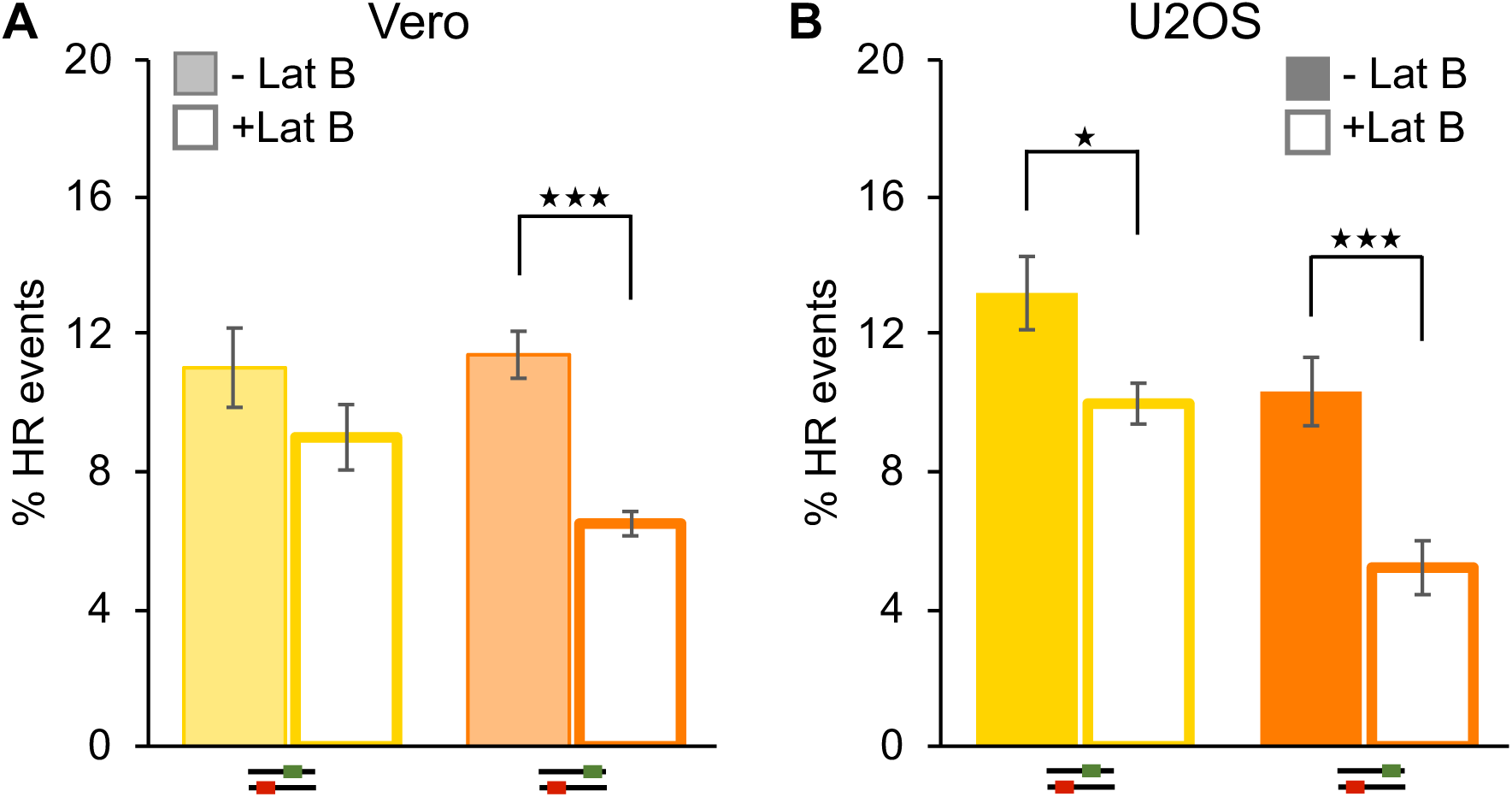
Inhibition of RC movement reduces the rate of HR recombination. Vero **(A)** or U2OS **(B)** cells were co-infected with the viral recombinants OK35 & OK25 (blue), OK35 & OK32 (yellow) and OK35 & OK26 (orange) in the presence (empty bar) or absence (full bar) of Latruncalin B. Progeny viruses from each infection condition were plated for individual plaques and colors were quantitated. The percent of dual color plaques out of the total fluorescent plaques were measured. The averages are shown from two individual plaque plating, from two technical repeats in two separate experiments. Error bars represent standard error of the means. *P < 0.05, ***P < 0.001; by t test.

### Triple-color FISH indicates that inter-genomic recombination occurs at edge of coalescing RCs

We speculated that if there are RCs that originate from a single recombinant genome and contain both tag sequences, there must be reciprocal recombinant RCs that contain none of the tags and are undetectable in our FISH assay (Figure 1C II). To determine whether these RCs exist, we designed an additional probe conjugated to a third fluorophore corresponding to the genomic HSV a’ sequence, a repetitive sequence found in four copies within the HSV-1 genome (56), This probe stains all HSV-1 viral DNA regardless of the tag sequence inserted. Both Vero and U2OS cells were co-infected with two isolates containing tags at different genomic loci, fixed and hybridized to all three probes as previously described. We inspected over 600 RCs from each cell type and found that 99.6% stained by the HSV non-specific probes reacted to either of the two specific probe sets (example in Figure 7). The existence of at least one tag sequence for all RCs suggests that RCs are not generated from recombinant genomes. This result further supports the hypothesis that recombination occurs at the edge of mature coalescing RCs at later stages of infection and is coupled with replication. Similarly, all coalescing areas between mature RCs respond to both the specific probe sets in contrast to our prediction that half of these areas should not respond to either specific probe (Figure 1C III). This finding suggests that these sites of recombination contain a mixture of genomes that contain either both tags or none of them. We therefore hypothesize that these sites of recombination do not arise from a single recombination event (since this would have led to the phenomenon of both tags or none) but rather from multiple events along the contact front of replicating RCs.

**Figure 7.**
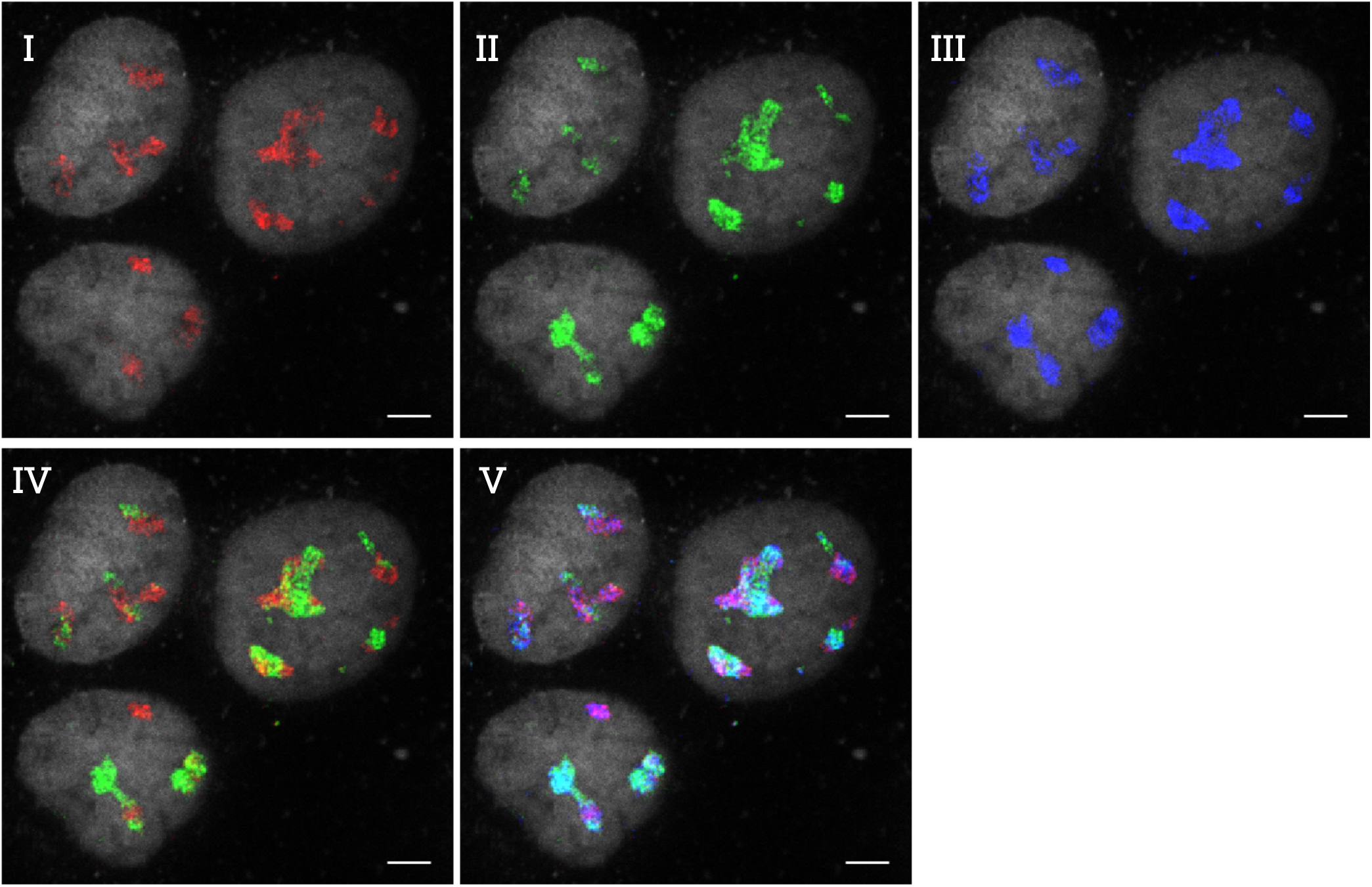
Replication compartments contain at least one tag sequence. Representative images of U2OS cells infected with two viral isolates containing tag sequences at different genomic loci (OK35 and OK32). The cells were hybridized with two probes conjugated to three distinct fluorophores. I. Detection of the OK35 tag sequence is shown in red. II. Detection of the OK32 tag sequence is shown in green. III. Detection of the viral ‘a’ sequence (detect HSV DNA non-specifically) is shown in blue. Overlays of two and three colours are shown in panels IV and V, respectively. DAPI is presented in gray. Scale bar is 5 µm.

## Discussion

Recombination among co-infecting herpesviruses is fundamentally important for understanding viral evolution and pathogenesis (5). It is also important to consider inter-genomic recombination when developing vectors for vaccination and oncolytic viral therapy, since evidence for recombination between a vaccine strain and a wild type strain has already been found in herpesviruses (57). Here, we developed a unique experimental system to identify the spatiotemporal constraints of inter-genomic recombination. Our results corroborate the hypothesis that each RC initiates from a single incoming genome. We visualized HR increasing in the overlapping areas between RCs, although the experimental system also detects overlapping signals unrelated to HR. We found that most viral inter-genomic recombination events that we detected occur at the edges of developing RCs where they coalesce with others. Inhibition of RCs movement reduced the rate of recombination, further corroborating that HR events occur after coalescencing of RCs. Finally, we suggest that areas in which overlapping RCs are detected result from multiple independent recombination events.

We have infected cells with different combinations of viral isolates: OK35 & OK25 where tags are inserted into the same genomic locus, OK35 & OK32 containing tags on separate genomic loci spaced approximately 72 Kbp apart, and OK35 & OK26 where they are spaced approximately 105 Kbp apart. In our analysis, we have not detected any significant differences between the two combinations of viruses tagged at separate genomic loci (i.e. OK35 & OK32 and OK35 & OK26). This supports the hypothesis that recombination events occur between replicating genomes that are in either branched or circular concatemeric states (7). Since HSV-1 maintains the 4 genome isomers at equivalent distributions (58), the 105Kbp distance between the markers is actually ~50Kbp in half of the concatemeric forms. This may explain the inability to distinguish differences in recombination rates when the markers are not closely linked, as previously suggested (7).

We observed four recognizable patterns of interactions between adjacent RCs: i. no mixing, ii. partial mixing, iii. one within another and iv. complete overlap (Figure 2 and 3). From our previous results with PRV (33), we expected that the lack of mixing of RCs would be the dominant interaction following co-infection with two viruses carrying tag sequences at the same location in the viral genomes (OK25 and OK35). We found that this readout is most commonly observed in this co-infection, although to our surprise the majority of RCs (~60%) showed some degree of mixing. Revisiting the images obtained during the Kobiler *et al.* paper (33) suggested that there is no major difference between the images from the two alphaherpesviruses co-infection. i.e the overlap between signals was also observed in the PRV experiments, but was attributed to image resolution and noise. We note that robust quantitative analysis of PRV images was not carried out.

The partial mixing of RCs was the most common interaction observed among the different co-infection conditions tested (Figure 2E). This is in part due to the categorization of all partial mixing interactions into one single pattern, regardless of the proportion of mixing. To overcome this problem, we developed an image analysis code that calculates the relative area of mixing. We found a significant increase in the relative area of mixing when co-infections were carried out with viruses in which HR can result in the mixing of the colors. These measurements were significant both at the single cell level (Figure 4A and B) and at the single RC level, indicating that a significant proportion of the mixing we observe is due to HR. On the other hand, even when HR cannot allow for signal mixing (i.e. OK25 and OK35 co-infection), we observed RCs and cells in which a high proportion of mixing occurred (Figure 4E and F). Similar findings were found in PRV co-infection assays (33). These results suggest that some color mixing can occur without recombining both tag sequences into the same genome. Alternatively they could reflect non-HR events during replication, which were previously observed both in HSV-1 replication (58) and in other herpesviruses (59). These explanations are not mutually exclusive, and both may contribute to the mixing of colors observed without HR.

The interaction in which one RC is fully mixed into a larger RC (Figure 2C) can be explained by multiple mechanisms. First, it could originate from a small RC that coalesces with a larger one. Second, the smaller RC could interact with two larger RCs that flank both sides of the small RC and eventually unite into a single larger RC. A third mechanism could involve the recently observed condensed viral genomes at the edge of RCs (34). Sekine *et al.* showed that at 3HPI several incoming genomes remain condensed at the periphery of an enlarging RC (34). These condensed genomes might be silent throughout the lytic infection or may serve as templates for recombination at later time points (which will result in our observed third interaction). The latter hypothesis fits our finding that this interaction is less frequent in U2OS (Figure 2E and 3D). Since U2OS cells lack ATRX and has compromised expression of other intrinsic immunity proteins (60–62), they are less likely to silence incoming genomes (52), and we speculate that they may have lower amounts of condensed genomes. Further experiments are required to distinguish between these different possibilities.

Our experimental system provides the opportunity to calculate accurately the number of functional RCs within a nucleus, since the separation by color increases our ability to differentiate between coalesced RCs. We were able to detect an increase in the average number of RCs detected in U2OS nuclei compared to Vero nuclei. This corroborates our previous finding that in U2OS cells on average a higher number of incoming viral genomes per cell initiate expression and replication compared to Vero cells (52). These observations also further support the emerging view that individual RCs initiate from single incoming genomes (33).

We define correlations between the nuclear area and the number of detectable RCs both in Vero and U2OS cells, similar to the correlation between cell size (as measured by flow cytometry) and the number of replicating genomes (54). Since our images represent snapshots during the infection process, we cannot distinguish between the possibilities that larger nuclei allow a larger number of incoming viral genomes to initiate replication or that the nuclei in which more genomes initiated replication expand faster. The use of new methods to visualise incoming genomes using click chemistry (34,35,63) should be able to resolve these two alternatives.

We note that U2OS cells are known to have larger nuclei and indeed have more RCs per cell than Vero cells. Similarly, HeLa cells that are relatively small cells with small nuclei have fewer incoming HSV-1 genomes that initiate replication (54). However, both U2OS and HeLa cells originated from tumors and carry multiple abnormalities that might regulate the number of herpes genomes that initiate replication. We are not certain that average nuclear size of a cell type is a main factor in determining the number of herpes genomes initiating replication among cell types.

Our analysis of the dual color FISH experiments led us to hypothesise that most viral inter-genomic HR events occur at the edge of the RCs and only after they coalesce. Further support for this hypothesis was obtained from the latrunculin experiment. Since viral RCs movement in the nucleus is dependent of actin filaments (31), the inhibition of actin polymerisation by the latrunculin B inhibitor should reduce RCs coalescence. Thus, the reduction in inter-genomic HR rates observed with latrunculin treatment can be attributed to a reduction in interaction between growing RCs. We note that when RCs enlarge, eventually, they will probably interact with each other without direct motion.

In mammalian nuclei, translocations between chromosomes, that can lead to tumorgenicity and other illnesses, are thought to occur among adjacent chromosomes (64,65). Our results support the requirement of proximity (contact first) among DNA molecules prior to recombination. Thus, maintaining genome integrity may require physical separation between DNA molecules. It will be interesting to determine what defines the location of viral replication compartments and the extent to which liquid phase separation (66) and other mechanisms regulate their mingling.

In conclusion, our results support a model in which incoming viral genomes establish RCs at distinct sites in the nucleus. The vast majority of RCs initiate from a single genome and only at later stages coalesce but maintain a detectable separation between RCs. These areas of interactions between the expanding RCs are predicted to be the site in which HR occurs in tight association with viral DNA replication. Our results suggest that in these sites, inter-genomic recombination is a frequent event, although not all areas of interactions between the RCs are supportive of close contact that can lead to recombination. Further research will shed light on the differences between these types of RC interactions.

## Funding

This work was supported by grants from the Israel Science foundation (grant #1387/14) to O.K., the EU CIG grant (FP7-2012-333653) to O.K., the National Institute of Health (NS082240) to M.D.W., the United States - Israel Binational Science Foundation (grant # 2015395) to O.K. and M.D.W., and a Marguerite Stolz Research Fellowship to O.K. E.T. is supported by the Buchman Scholarship and A.Z. was supported by the National Institute of Health Grant to Gaudenz Danuser (P01 GM103723). The funders had no role in study design, data collection and analysis, decision to publish, or preparation of the manuscript.

## Acknowledgement

We thank Eitan Erez Zahavi, Ariel Ionescu and Eran Perlson for advice and valuable support and all Kobiler lab members for their comments.

---

**Supplementary table 1. Accumulated data from image analysis program outputs.** Data from 3 experiments for each cell type was collected. Each line represents individual cell. For each cell, the following parameters are listed: the number of RCs in that cell, the number of colocalization events, the area of the nucleus, the total area of all RC, the total area of colocalization events, the adjusted RC area (total area of the RCs minus the area of colocalization which removes measuring the area of colocalization in both colors) and the ratio between area of colocalization to adjusted RC area.

## References

1. Muller, H.J. (1964) The Relation of Recombination to Mutational Advance. Mutat Res, 106, 2–9.

2. Felsenstein, J. (1974) The evolutionary advantage of recombination. Genetics, 78, 737–756.

3. Sanjuan, R. and Domingo-Calap, P. (2016) Mechanisms of viral mutation. Cell Mol Life Sci, 73, 4433–4448.

4. Holmes, E.C. (2008) Evolutionary history and phylogeography of human viruses. Annu Rev Microbiol, 62, 307–328.

5. Renner, D.W. and Szpara, M.L. (2017) The Impacts of Genome-wide Analyses on our Understanding of Human Herpesvirus Diversity and Evolution. J Virol.

6. Brown, S.M., Ritchie, D.A. and Subak-Sharpe, J.H. (1973) Genetic studies with herpes simplex virus type 1. The isolation of temperature-sensitive mutants, their arrangement into complementation groups and recombination analysis leading to a linkage map. J Gen Virol, 18, 329–346.

7. Honess, R.W., Buchan, A., Halliburton, I.W. and Watson, D.H. (1980) Recombination and linkage between structural and regulatory genes of herpes simplex virus type 1: study of the functional organization of the genome. J Virol, 34, 716–742.

8. Law, G.A., Herr, A.E., Cwick, J.P. and Taylor, M.P. (2018) A New Approach to Assessing HSV-1 Recombination during Intercellular Spread. Viruses, 10.

9. Lee, K., Kolb, A.W., Sverchkov, Y., Cuellar, J.A., Craven, M. and Brandt, C.R. (2015) Recombination Analysis of Herpes Simplex Virus 1 Reveals a Bias toward GC Content and the Inverted Repeat Regions. J Virol, 89, 7214–7223.

10. Morse, L.S., Buchman, T.G., Roizman, B. and Schaffer, P.A. (1977) Anatomy of herpes simplex virus DNA. IX. Apparent exclusion of some parental DNA arrangements in the generation of intertypic (HSV-1 X HSV-2) recombinants. J Virol, 24, 231–248.

11. Schaffer, P.A., Aron, G.M., Biswal, N. and Benyesh-Melnick, M. (1973) Temperature-sensitive mutants of herpes simplex virus type 1: isolation, complementation and partial characterization. Virology, 52, 57–71.

12. Tang, K.W., Norberg, P., Holmudden, M., Elias, P. and Liljeqvist, J.A. (2014) Rad51 and Rad52 are involved in homologous recombination of replicating herpes simplex virus DNA. PLoS One, 9, e111584.

13. Szpara, M.L., Gatherer, D., Ochoa, A., Greenbaum, B., Dolan, A., Bowden, R.J., Enquist, L.W., Legendre, M. and Davison, A.J. (2014) Evolution and diversity in human herpes simplex virus genomes. J Virol, 88, 1209–1227.

14. Kolb, A.W., Ane, C. and Brandt, C.R. (2013) Using HSV-1 genome phylogenetics to track past human migrations. PLoS One, 8, e76267.

15. Bowden, R., Sakaoka, H., Donnelly, P. and Ward, R. (2004) High recombination rate in herpes simplex virus type 1 natural populations suggests significant co-infection. Infect Genet Evol, 4, 115–123.

16. Kobiler, O., Drayman, N., Butin-Israeli, V. and Oppenheim, A. (2012) Virus strategies for passing the nuclear envelope barrier. Nucleus, 3, 526–539.

17. Burkham, J., Coen, D.M. and Weller, S.K. (1998) ND10 protein PML is recruited to herpes simplex virus type 1 prereplicative sites and replication compartments in the presence of viral DNA polymerase. J Virol, 72, 10100–10107.

18. Dembowski, J.A. and DeLuca, N.A. (2015) Selective recruitment of nuclear factors to productively replicating herpes simplex virus genomes. PLoS Pathog, 11, e1004939.

19. Lilley, C.E., Chaurushiya, M.S., Boutell, C., Everett, R.D. and Weitzman, M.D. (2011) The intrinsic antiviral defense to incoming HSV-1 genomes includes specific DNA repair proteins and is counteracted by the viral protein ICP0. PLoS Pathog, 7, e1002084.

20. Liptak, L.M., Uprichard, S.L. and Knipe, D.M. (1996) Functional order of assembly of herpes simplex virus DNA replication proteins into prereplicative site structures. J Virol, 70, 1759–1767.

21. Mohni, K.N., Livingston, C.M., Cortez, D. and Weller, S.K. (2010) ATR and ATRIP are recruited to herpes simplex virus type 1 replication compartments even though ATR signaling is disabled. J Virol, 84, 12152–12164.

22. Reyes, E.D., Kulej, K., Pancholi, N.J., Akhtar, L.N., Avgousti, D.C., Kim, E.T., Bricker, D.K., Spruce, L.A., Koniski, S.A., Seeholzer, S.H. et al. (2017) Identifying Host Factors Associated with DNA Replicated During Virus Infection. Mol Cell Proteomics, 16, 2079–2097.

23. Taylor, T.J. and Knipe, D.M. (2004) Proteomics of herpes simplex virus replication compartments: association of cellular DNA replication, repair, recombination, and chromatin remodeling proteins with ICP8. J Virol, 78, 5856–5866.

24. Vogel, J.L. and Kristie, T.M. (2013) The dynamics of HCF-1 modulation of herpes simplex virus chromatin during initiation of infection. Viruses, 5, 1272–1291.

25. Zhong, L. and Hayward, G.S. (1997) Assembly of complete, functionally active herpes simplex virus DNA replication compartments and recruitment of associated viral and cellular proteins in transient cotransfection assays. J Virol, 71, 3146–3160.

26. Roizman, B. and Zhou, G. (2015) The 3 facets of regulation of herpes simplex virus gene expression: A critical inquiry. Virology, 479–480, 562-567.

27. Quinlan, M.P., Chen, L.B. and Knipe, D.M. (1984) The intranuclear location of a herpes simplex virus DNA-binding protein is determined by the status of viral DNA replication. Cell, 36, 857–868.

28. de Bruyn, Kops A. and Knipe, D.M. (1988) Formation of DNA replication structures in herpes virus-infected cells requires a viral DNA binding protein. Cell, 55, 857–868.

29. Lukonis, C.J. and Weller, S.K. (1996) Characterization of nuclear structures in cells infected with herpes simplex virus type 1 in the absence of viral DNA replication. J Virol, 70, 1751–1758.

30. Taylor, T.J., McNamee, E.E., Day, C. and Knipe, D.M. (2003) Herpes simplex virus replication compartments can form by coalescence of smaller compartments. Virology, 309, 232–247.

31. Chang, L., Godinez, W.J., Kim, I.H., Tektonidis, M., de Lanerolle, P., Eils, R., Rohr, K. and Knipe, D.M. (2011) Herpesviral replication compartments move and coalesce at nuclear speckles to enhance export of viral late mRNA. Proc Natl Acad Sci U S A, 108, E136–144.

32. Sourvinos, G. and Everett, R.D. (2002) Visualization of parental HSV-1 genomes and replication compartments in association with ND10 in live infected cells. The EMBO journal, 21, 4989–4997.

33. Kobiler, O., Brodersen, P., Taylor, M.P., Ludmir, E.B. and Enquist, L.W. (2011) Herpesvirus replication compartments originate with single incoming viral genomes. MBio, 2.

34. Sekine, E., Schmidt, N., Gaboriau, D. and O’Hare, P. (2017) Spatiotemporal dynamics of HSV genome nuclear entry and compaction state transitions using bioorthogonal chemistry and super-resolution microscopy. PLoS Pathog, 13, e1006721.

35. Dembowski, J.A. and DeLuca, N.A. (2018) Temporal Viral Genome-Protein Interactions Define Distinct Stages of Productive Herpesviral Infection. MBio, 9.

36. Reuven, N.B., Staire, A.E., Myers, R.S. and Weller, S.K. (2003) The herpes simplex virus type 1 alkaline nuclease and single-stranded DNA binding protein mediate strand exchange in vitro. J Virol, 77, 7425–7433.

37. Schumacher, A.J., Mohni, K.N., Kan, Y.N., Hendrickson, E.A., Stark, J.M. and Weller, S.K. (2012) The HSV-1 Exonuclease, UL12, Stimulates Recombination by a Single Strand Annealing Mechanism. Plos Pathogens, 8.

38. Weller, S.K. and Coen, D.M. (2012) Herpes Simplex Viruses: Mechanisms of DNA Replication. Csh Perspect Biol, 4.

39. Advani, S.J., Weichselbaum, R.R. and Roizman, B. (2003) Herpes simplex virus 1 activates cdc2 to recruit topoisomerase II alpha for post-DNA synthesis expression of late genes. Proc Natl Acad Sci U S A, 100, 4825–4830.

40. Edwards, T.G., Bloom, D.C. and Fisher, C. (2018) The ATM and Rad3-Related (ATR) Protein Kinase Pathway Is Activated by Herpes Simplex Virus 1 and Required for Efficient Viral Replication. J Virol, 92.

41. Lilley, C.E., Carson, C.T., Muotri, A.R., Gage, F.H. and Weitzman, M.D. (2005) DNA repair proteins affect the lifecycle of herpes simplex virus 1. Proc Natl Acad Sci U S A, 102, 5844–5849.

42. Mohni, K.N., Dee, A.R., Smith, S., Schumacher, A.J. and Weller, S.K. (2013) Efficient herpes simplex virus 1 replication requires cellular ATR pathway proteins. J Virol, 87, 531–542.

43. De Chiara, G., Racaniello, M., Mollinari, C., Marcocci, M.E., Aversa, G., Cardinale, A., Giovanetti, A., Garaci, E., Palamara, A.T. and Merlo, D. (2016) Herpes Simplex Virus-Type1 (HSV-1) Impairs DNA Repair in Cortical Neurons. Front Aging Neurosci, 8, 242.

44. Parkinson, J., Lees-Miller, S.P. and Everett, R.D. (1999) Herpes simplex virus type 1 immediate-early protein vmw110 induces the proteasome-dependent degradation of the catalytic subunit of DNA-dependent protein kinase. J Virol, 73, 650–657.

45. Smith, S., Reuven, N., Mohni, K.N., Schumacher, A.J. and Weller, S.K. (2014) Structure of the herpes simplex virus 1 genome: manipulation of nicks and gaps can abrogate infectivity and alter the cellular DNA damage response. J Virol, 88, 10146–10156.

46. Smith, S. and Weller, S.K. (2015) HSV-I and the cellular DNA damage response. Future Virol, 10, 383–397.

47. Trigg, B.J., Lauer, K.B., Fernandes Dos Santos, P., Coleman, H., Balmus, G., Mansur, D.S. and Ferguson, B.J. (2017) The Non-Homologous End Joining Protein PAXX Acts to Restrict HSV-1 Infection. Viruses, 9.

48. Wilkinson, D.E. and Weller, S.K. (2006) Herpes simplex virus type I disrupts the ATR-dependent DNA-damage response during lytic infection. J Cell Sci, 119, 2695–2703.

49. Taylor, M.P., Kobiler, O. and Enquist, L.W. (2012) Alphaherpesvirus axon-to-cell spread involves limited virion transmission. Proc Natl Acad Sci U S A, 109, 17046–17051.

50. Cremer, M., Grasser, F., Lanctot, C., Muller, S., Neusser, M., Zinner, R., Solovei, I. and Cremer, T. (2008) Multicolor 3D fluorescence in situ hybridization for imaging interphase chromosomes. Methods Mol Biol, 463, 205–239.

51. Otsu, N. (1979) Threshold Selection Method from Gray-Level Histograms. Ieee T Syst Man Cyb, 9, 62–66.

52. Shapira, L., Ralph, M., Tomer, E., Cohen, S. and Kobiler, O. (2016) Histone Deacetylase Inhibitors Reduce the Number of Herpes Simplex Virus-1 Genomes Initiating Expression in Individual Cells. Front Microbiol, 7, 1970.

53. Li, Z., Fang, C., Su, Y., Liu, H., Lang, F., Li, X., Chen, G., Lu, D. and Zhou, J. (2016) Visualizing the replicating HSV-1 virus using STED super-resolution microscopy. Virol J, 13, 65.

54. Cohen, E.M. and Kobiler, O. (2016) Gene Expression Correlates with the Number of Herpes Viral Genomes Initiating Infection in Single Cells. PLoS Pathog, 12, e1006082.

55. Drayman, N., Karin, O., Mayo, A., Danon, T., Shapira, L., Rafael, D., Zimmer, A., Bren, A., Kobiler, O. and Alon, U. (2017) Dynamic Proteomics of Herpes Simplex Virus Infection. MBio, 8.

56. Hayward, G.S., Jacob, R.J., Wadsworth, S.C. and Roizman, B. (1975) Anatomy of herpes simplex virus DNA: evidence for four populations of molecules that differ in the relative orientations of their long and short components. Proc Natl Acad Sci U S A, 72, 4243–4247.

57. Lee, S.W., Markham, P.F., Coppo, M.J., Legione, A.R., Markham, J.F., Noormohammadi, A.H., Browning, G.F., Ficorilli, N., Hartley, C.A. and Devlin, J.M. (2012) Attenuated vaccines can recombine to form virulent field viruses. Science, 337, 188.

58. Mahiet, C., Ergani, A., Huot, N., Alende, N., Azough, A., Salvaire, F., Bensimon, A., Conseiller, E., Wain-Hobson, S., Labetoulle, M. et al. (2012) Structural variability of the herpes simplex virus 1 genome in vitro and in vivo. J Virol, 86, 8592–8601.

59. Rijsewijk, F.A., Verschuren, S.B., Madic, J., Ruuls, R.C., Renaud, P. and van Oirschot, J.T. (1999) Spontaneous BHV1 recombinants in which the gI/gE/US9 region is replaced by a duplication/inversion of the US1.5/US2 region. Arch Virol, 144, 1527–1537.

60. Heaphy, C.M., de Wilde, R.F., Jiao, Y., Klein, A.P., Edil, B.H., Shi, C., Bettegowda, C., Rodriguez, F.J., Eberhart, C.G., Hebbar, S. et al. (2011) Altered telomeres in tumors with ATRX and DAXX mutations. Science, 333, 425.

61. Hancock, M.H., Corcoran, J.A. and Smiley, J.R. (2006) Herpes simplex virus regulatory proteins VP16 and ICP0 counteract an innate intranuclear barrier to viral gene expression. Virology, 352, 237–252.

62. Yao, F. and Schaffer, P.A. (1995) An activity specified by the osteosarcoma line U2OS can substitute functionally for ICP0, a major regulatory protein of herpes simplex virus type 1. J Virol, 69, 6249–6258.

63. Alandijany, T., Roberts, A.P.E., Conn, K.L., Loney, C., McFarlane, S., Orr, A. and Boutell, C. (2018) Distinct temporal roles for the promyelocytic leukaemia (PML) protein in the sequential regulation of intracellular host immunity to HSV-1 infection. PLoS Pathog, 14, e1006769.

64. McCord, R.P. and Balajee, A. (2018) 3D Genome Organization Influences the Chromosome Translocation Pattern. Adv Exp Med Biol, 1044, 113–133.

65. Misteli, T. and Soutoglou, E. (2009) The emerging role of nuclear architecture in DNA repair and genome maintenance. Nat Rev Mol Cell Biol, 10, 243–254.

66. McSwiggen, D.T., Hansen, A.S., Marie-Nelly, H., Teves, S., Heckert, A., Dugast-Darzacq, C., Hao, Y., Umemoto, K.K., Tjian, R. and Darzacq, X. (2018) Transient DNA Binding Induces RNA Polymerase II Compartmentalization During Herpesviral Infection Distinct From Phase Separation. bioRxiv.

